# Gibberellins promote polar auxin transport to regulate stem cell fate decisions in cambium

**DOI:** 10.1101/2022.07.15.500224

**Authors:** Riikka Mäkilä, Brecht Wybouw, Ondrej Smetana, Leo Vainio, Anna Solé-Gil, Munan Lyu, Lingling Ye, Xin Wang, Riccardo Siligato, Mark Kubo Jenness, Angus S. Murphy, Ari Pekka Mähönen

## Abstract

Vascular cambium contains bifacial stem cells, which produce secondary xylem to one side and secondary phloem to the other. However, how these fate decisions are regulated is unknown. Here, we show that the positioning of an auxin signalling maximum within the cambium determines the fate of stem cell daughters. The position is modulated by gibberellin-regulated, PIN1-dependent polar auxin transport. Gibberellin treatment broadens auxin maximum from the xylem side of the cambium towards the phloem. As a result, xylem-side stem cell daughter preferentially differentiates into xylem, while phloem-side daughter retains stem cell identity. Occasionally, this broadening leads to direct specification of both daughters as xylem, and consequently, adjacent phloem-identity cell reverts to being stem cell. Conversely, reduced gibberellin levels favour specification of phloem-side stem cell daughter as phloem. Together, our data provide a mechanism by which gibberellin regulates the ratio of xylem and phloem production.

## Main

Vascular cambium is responsible for the lateral (secondary) growth of plant stems and roots. This process is particularly prevalent in tree species but also occurs in non-woody species like *Arabidopsis thaliana*^1^. The vascular cambium consists of meristematic cells that undergo periclinal cell divisions (that is, cell divisions parallel to the surface of the organ)^2^. Cambium cells that leave the meristem ultimately differentiate into parenchymatic or conductive cells, with secondary xylem being produced inwards and secondary phloem outwards^3^ (**Extended Data Fig. 1a**). Recent lineage-tracing studies showed that a subset of cambial cells act as bifacial stem cells, since a single cambial cell is capable of producing both xylem and phloem^4–6^.

A major regulator of cambium development is the phytohormone auxin^4,7,8^. Mutations in genes encoding components of auxin signalling including those associated with perception and polar transport of the hormone cause defects in cambium development^4,9^, vascular patterning^4,9–11^, leaf venation^12^, xylem and phloem formation *in planta*^4,13,14^, in tissue culture^15^ and during vascular regeneration^16^. Recently, we showed that ectopic clones with high levels of auxin signalling force non-xylem cells to differentiate into secondary xylem vessels, while cells adjacent to such clones divide periclinally and gain expression of cambial markers^4^. The ectopic clone thus behaves as an organizer that causes adjacent cells to specify as vascular cambium stem cell-like cells. In agreement with this, an auxin maximum is normally located on the xylem side of the vascular cambium, and stem cell divisions occur adjacent to this maximum^4^. These data raise the question whether the location of the auxin maximum within the cambium has a role in stem cell fate decisions.

Other phytohormones also influence cambium development alongside auxin^8^. For example, gibberellins (or gibberellic acid, GA) promote secondary xylem production in both *Arabidopsis*^17,18^ and poplar^18–20^. In *Arabidopsis*, this occurs during flowering, when GA levels rise^17^. Recently, *AUXIN RESPONSE FACTORs 6* (*arf6*) and *ARF8* have been shown to mediate auxin-dependent xylem production that is downstream of GA^21^. Interactions between auxin and GA also occur in other biological processes. For example, in *Arabidopsis* roots, GA directly promotes abundance of PIN polar auxin transporters in the root meristem, thus regulating polar auxin transport (PAT)^22^.

In this work, we show that GA promotes PIN1-dependent PAT in *Arabidopsis thaliana* roots. This results in an expanded auxin signalling maximum within the root vascular cambium, which forces cambial stem cell daughters to preferentially specify as xylem cells. Our data show how GA influence the position of the auxin maximum in cambium, therefore determining stem cell fate decisions between xylem and phloem.

### GA regulates stem cell fate decisions

Previously, GA has been shown to increase xylem formation in *Arabidopsis* hypocotyls during flowering^17^. In order to understand the role of GA on cambial growth dynamics, we analysed GA’s effect in *Arabidopsis* roots at a cellular resolution. To reach that goal, we analysed roots during the early stages of secondary growth, when cell division and differentiation dynamics are easier to follow. At these stages, only two types of xylem cells are produced: secondary xylem vessels and xylem parenchyma (**Extended Data Fig. 1a**). Secondary xylem vessels expand radially and deposit a thick secondary cell wall before fully differentiating into hollow, water-conducting vessels, while xylem parenchyma remain in an undifferentiated state. As expected, GA treatment in young roots resulted in an increased number of both secondary xylem vessels and xylem parenchyma, and the increase was equal in both cell types (**Fig. 1a,b,c**; **Extended Data Fig. 1b**). In addition, secondary xylem vessel expansion increased as a result of GA treatment (**Fig. 1a,d**). In contrast to plants treated with GA, a mutant deficient in GA biosynthesis, *ga1*^23^, had a reduced number of xylem vessels and parenchymatic cells (**Fig. 1a,b,c**). Additionally, the xylem vessel area was reduced (**Fig. 1a,d**). All of these phenotypes were rescued by GA treatment (**Fig. 1a,b,c,d**). Altogether, these data show that GA promotes the production of both xylem vessels and parenchyma during the early stages of secondary growth in roots.

**Figure 1.**
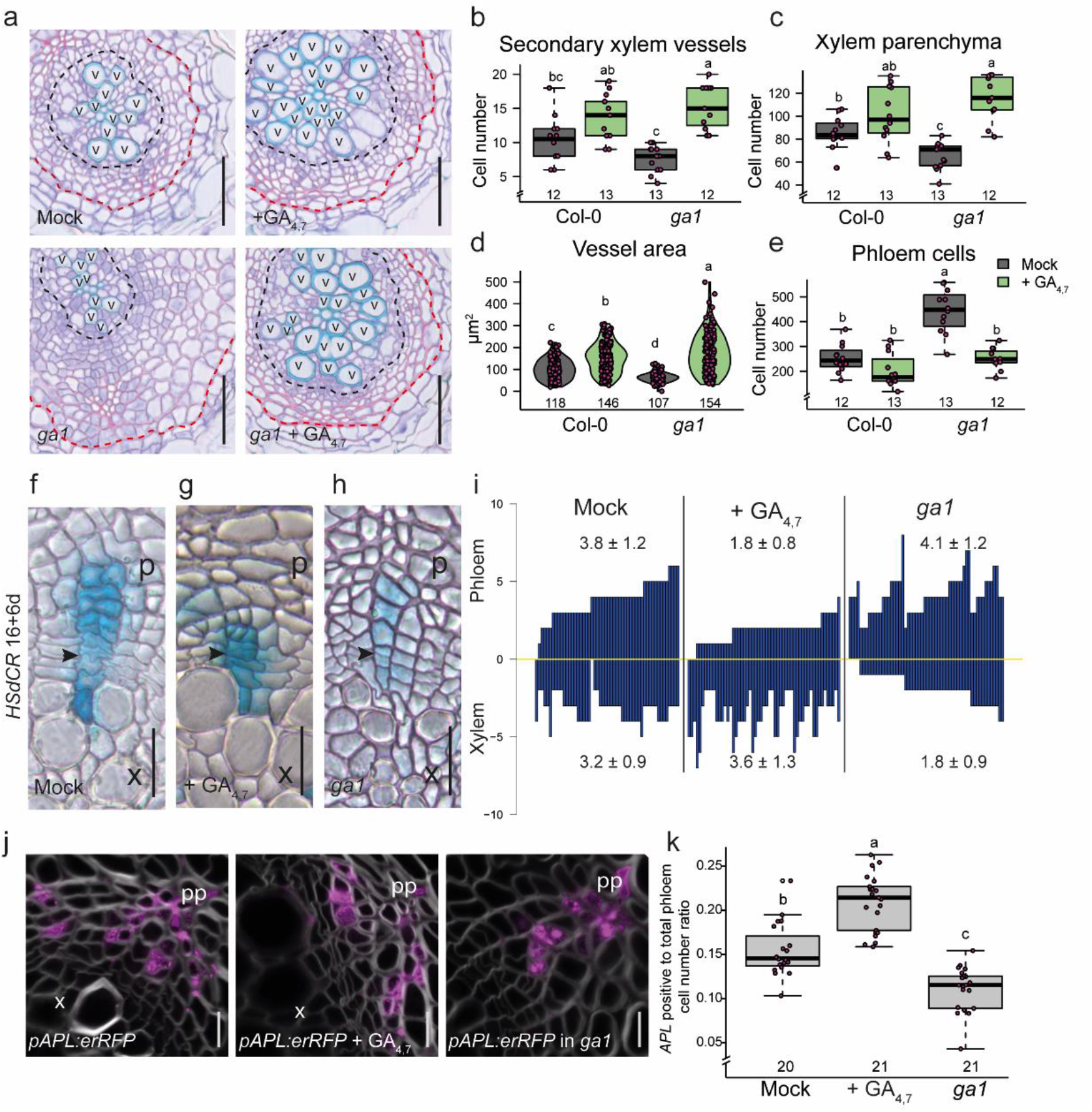
GA induces secondary xylem proliferation and vessel expansion. (**a**) Root cross-sections after a 10-day GA treatment in 4-day old Col-0 and *ga1* plants. Black dotted lines indicate the most recent cell divisions. Red dotted lines mark the border between the phloem parenchyma cells and the periderm. (**b-e**) Quantifications of secondary vessel (**b**) and xylem parenchyma cell numbers (**c**), individual secondary vessel area (**d**), and total phloem cell number (**e**) in 14-day old seedlings. (**f-i**) Lineage tracing in active cambium with GUS stained sectors (blue) originating from a single recombination event. Recombination was induced in 16-day old seedlings, after which the seedlings (Col-0 in **f,g** and *ga1* in **h**) were grown for an additional 6 days under mock (**f,h**) or GA_4,7_ conditions (**g**). Black arrowheads indicate the most recent cell divisions in the sectors, where the thinnest cell wall was observed. (**i**) GUS sectors (bars) plotted relative to the position of the thinnest cell wall (yellow line) in each sector. Values above and below the bars indicate average number of phloem and xylem cells (±SD), respectively, within the sectors. Some of the sectors, especially after GA treatment, ended on xylem vessels, which are dead and thus cannot be observed with GUS staining. Therefore, the length of these sectors towards the xylem is an underestimation of the actual length. (**j**) Confocal cross sections of *pAPL:erRFP* after an 11-day GA treatment in 4-day old plants (except *pAPL:erRFP* in the *ga1* mutant background, which was grown 15 days in Mock). The *APL* reporter marks conductive phloem cells. (**k**) The ratio of cells expressing *APL* versus all phloem cells in **j**. In **b,c,e** and **k**, the boxes in the box and whisker plots show the median and interquartile range, and the whiskers show the total range. Individual data points are plotted as purple dots. In the violin plots in **d**, the white dot shows the median and the thick line the interquartile range. The thinner line represents the rest of the distribution. Each side of the line is a kernel density estimation that shows the distribution shape of the data. Individual data points are plotted as purple dots. Numbers in **b-e** and **k** indicate number of samples. Two-way ANOVA with Tukey’s post hoc test in **b-e** and **k**. Letters indicate a significant difference, *P* < 0.05. Scale bars are 50 μm (**a**), 20 μm (**f-h**) or 10 μm (**j**). “p” = phloem, “pp” = primary phloem pole, “x” = xylem “v” = secondary xylem vessel. All experiments were repeated three times.

To investigate the mechanism causing the observed changes in xylem cell number, we looked for alterations in the cambium growth dynamics. We used a previously established a heat shock inducible CRE–*lox* based lineage-tracing system (*HSdCR*)^4^ which allows the production of single-cell clones within a population of dividing cells, including cambium. This enabled us to monitor the cambium growth dynamics over time. Under normal growth conditions, lineages are derived from a single recombination event in one stem cell and span towards both the xylem and phloem side in an almost equal manner (**Fig. 1f,i**). This indicates that bifacial stem cell divisions normally provide an equal number of new xylem and phloem cells. Under GA-treated conditions, clone cell lineages show an unequal distribution (**Fig. 1g,i**), with a preference towards the xylem side, while lineages in the *ga1* mutant background preferably span towards the phloem (**Fig. 1h,i**). We did not observe proliferating sectors exiting the cambium and entering differentiating tissue in any of the conditions (**Fig. 1i**). These data indicate that GA regulates stem cell fate decisions during cambium proliferation rather than specifically regulating xylem or phloem proliferation.

### Dual function of GA on phloem formation

Previous histological studies in hypocotyl^21^ and our lineage-tracing results in root (Fig. 1i) show that GA inhibits phloem production. Next, we tested whether GA affects the production of different phloem cell types. Phloem consists of conductive cells known as sieve elements, together with their companion cells and phloem parenchyma (**Extended Data Fig. 1a**). In agreement with the lineage-tracing results, total phloem cell numbers were decreased in GA-treated roots and increased in the *ga1* mutant background (**Fig. 1a,e**). Next, we used the conductive phloem cell specific marker *ALTERED PHLOEM DEVELOPMENT* (*APL*)^24^ to determine whether GA affects the number of conductive phloem cells. We observed that the ratio of APL-positive cells to total phloem cells was increased after GA treatment and decreased in *ga1* (**Fig. 1j,k**). Thus, with excess GA, plants produce more conductive phloem, and with limited GA, they instead produce parenchymatic cells. Similar results were observed when quantifying the number of sieve elements by safranin staining^25^; the number of sieve elements was decreased in *ga1* (**Extended Data Fig. 1f,g**). These results seem counterintuitive compared to the lineage tracing and total phloem number results, where the reverse tendency was observed. We therefore analysed the overall expression pattern of *APL* in more detail. In *ga1*, *APL* expression showed that phloem differentiation is more focused around the primary phloem pole regions and is situated further away from the dividing stem cells than in normal conditions (**Extended Data Fig. 1c,d,e**). In contrast, after GA treatment, plants show broader *APL* expression, with phloem differentiation occurring slightly closer to the dividing stem cells **(Extended Data Fig. 1d,e**). These data indicate that GA inhibits a phloem fate decision by cambial stem cells; however, those few cells that do specify as phloem will preferentially differentiate as conductive phloem.

### GA signalling is required in the early xylem domain

Next, we wondered where and how GA affects cambium growth dynamics. First, we aimed to understand which tissue types GA signalling operates in during secondary growth. DELLA proteins act as repressors of GA signalling, and they are rapidly degraded in the presence of GA^26^. Mutations in one of the DELLA genes, *REPRESSOR OF GA* (*RGA*)^27^, result in increased xylem area^21^ within the hypocotyl and could therefore also have an effect in root secondary growth. Indeed, we found that *pRGA:GFP-RGA*^27^ showed broad expression in the root cambium, appearing in both the xylem and the phloem (**Fig. 2a)**, and a 6 h GA application led to degradation of the *pRGA:GFP-RGA* signal in all cell types (**Fig. 2a**), indicating that the GA signalling components are broadly present in secondary tissues.

**Figure 2.**
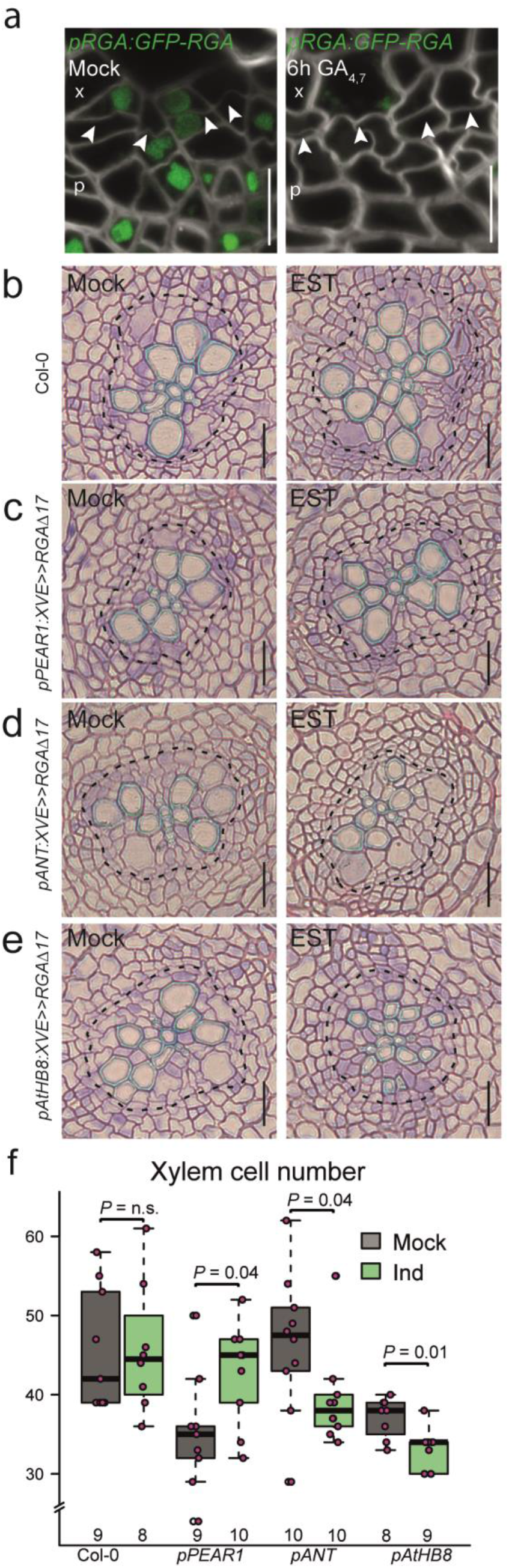
GA signalling on the xylem side of the cambium is required to promote secondary xylem formation. (**a**) Expression pattern of *pRGA:GFP-RGA* in the root cambium of 14-day old seedlings after a 6h mock or GA4,7 treatment. White arrows indicate the most recent divisions. (**b-e**) Root cross-sections after a 6-day induction in 4-day old seedlings of Col-0 (**b**) or with mutated RGAΔ17 expressed in the early phloem cell (*pPEAR1:XVE>>RGAΔ17*) (**c**), the stem cells (*pANT:XVE>>RGAΔ17*) (**d**), or the early xylem (*pATHB8:XVE>>RGAΔ17*) (**e**). Black dotted lines indicate the most recent divisions. (**f**) Quantification of the total xylem cell number (cells within the most recent cell divisions) in panels **b-e**. Scale bars are 10 μm (**a**) or 20 μm (**b-e**). Significant differences based on a two-tailed Wilcoxon-test are indicated. Numbers in **f** indicate number of samples. The boxes in the box and whisker plots show the median and the interquartile range, and the whiskers show the total range. Individual data points are plotted as purple dots. “p” = phloem, “x” = xylem. All experiments were repeated three times.

Deletion of 17 amino acids within the DELLA domain of RGA (RGAΔ17) results in the formation of a dominant, non-degradable version of the protein^28^. By driving this dominant inhibitor of GA signalling under three different cell type-specific inducible promoters, we investigated where GA signalling is required for its effect on cambium development. Inhibition of GA signalling under the promoter of the early phloem gene *PHLOEM-EARLY-DOF 1* (*PEAR1*)^29^ did not inhibit xylem production; unexpectedly, it led to an increase in xylem cell number (**Fig. 2b,c,f**). However, RGAΔ17 induction under the promoter of the stem cell gene *AINTEGUNMENTA (ANT*)^4^, and especially under the promoter of the early xylem gene *HOMEOBOX GENE 8 (AtHB8*)^4^ significantly reduced xylem production (**Fig. 2b,d-f**), with the strongest lines resembling the *ga1* mutant phenotype (**Fig. 1a** and **Fig. 2e**). These data indicate that GA signalling in the stem cells and in early xylem is required for its role in promoting xylem production. This is also in accordance with measured bioactive GA gradients within poplar stems^20^, which show a GA maximum in the developing xylem.

### GA regulates the width of the auxin response gradient to promote xylem formation

Earlier clonal activation studies have shown that a local auxin maximum drives xylem formation and promotes cambial cell divisions non-cell autonomously^4^. As GA’s effect on xylem proliferation is the strongest in the early xylem cells, where the local auxin signalling maximum is located, we investigated whether GA could regulate the position of this maximum. Using a new RFP-based version of the auxin response reporter, *DR5v2*^30^ (see Methods), we observed expression on the xylem side of cambium (**Fig. 3a**), matching which cells show the highest levels of auxin signalling in secondary tissues^4^. Recent stem cell divisions are identifiable by the appearance of thin cell walls within the cambium (arrowheads in **Fig. 3a**). We marked the phloem-side stem cell daughter as 1 and the xylem-side daughter as −1 (**Fig. 3a,b**). In wild type plants, *DR5v2* expression often reaches the xylem-side stem cell daughter (−1) and even reached the cell in position −2, but it was rarely seen in the phloem-side daughter. In *ga1*, a smaller proportion of stem cell daughters showed *DR5v2* expression (expression in positions 1 or −1 was seen in 29% of *ga1* roots and 48% of Col-0 roots, **Fig. 3b, Extended Data Fig. 2a**). A 24 h GA treatment was not sufficient to cause changes in *DR5v2* expression (**Extended data Fig. 2b-d**). However, after 48 h, a higher proportion of the stem cell daughters expressed *DR5v2* than in mock controls (57% in positions −1 and 1 in *ga1* and 83% in Col-0) (**Fig. 3a,b,c**). Altogether, these GA manipulation studies show that GA regulates the position of the auxin signalling maximum within cambium.

**Figure 3.**
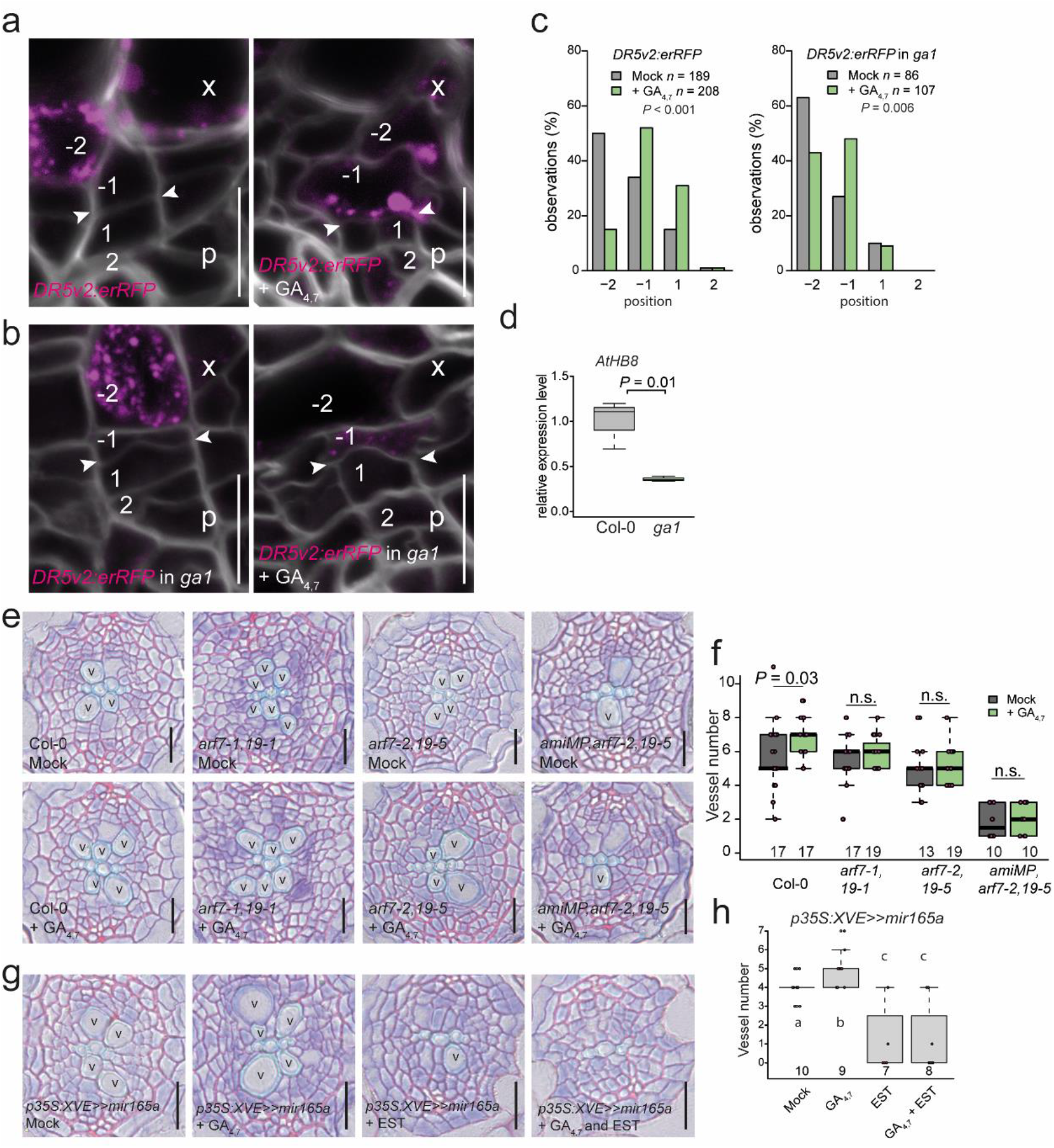
Auxin is required for GA to affect xylem development. Expression of *DR5v2:erRFP* in the root cambium after a 48 h GA treatment in 14-day old seedlings of wild type Col-0 (**a**) and *ga1* (**b**). White arrowheads indicate the most recent cell divisions. The numbers “-2”, “-1”, “1” and “2” indicate the relative position of the cells in respect to the most recent cell division, with negative values towards the xylem and positive towards the phloem. (**c**) Counts of the position in the cambium at which the *DR5v2:erRFP* gradient ends. Cellular positions on the x-axis correspond with the cellular position in panel **a** & **b**, and n refers to the total number of observations. (**d**) qRT-PCR analysis of the *AtHB8* expression level in wild type and *ga1* backgrounds. (**e**) Root cross-sections after a 6-day GA treatment in 4-day old seedlings of Col-0, *arf7,arf19*, and *amiMP,arf7,arf19*. (**f**) Quantification of the number of secondary xylem vessels in plants shown in panel **e**. (**g**) Root cross-sections after a 6-day induction and GA treatment in 4-day old seedlings of *p35S::XVE>>mir165a* seedlings. (**h**) Quantification of the number of secondary xylem vessels in plants shown in panel **g**. Chi-squared test in **c**; two-tailed t-test in **d,f**; two-way ANOVA with Tukey’s post hoc test in **h**. The boxes in the box and whisker plots show the median and the interquartile range, and the whiskers show the total range. Individual data points are plotted as purple dots. Numbers in **f** and **h** indicate number of samples. Letters indicate a significant difference, *P* < 0.05. “p”= phloem, “x”= xylem, “v”= secondary xylem vessels, n refers to the total number of observations. Scale bars are 10 μm (**a,b**) or 20 μm (**e,g**). All experiments were repeated three times.

Since auxin drives xylem vessel formation^4^, this GA-induced broadened auxin response gradient could explain how GA promotes vessel production (**Fig. 1a,b**). To the test this, we investigated whether auxin signalling is required for the effect of GA on xylem production in the root cambium. Previously, we have shown that auxin signalling in the *Arabidopsis* root cambium acts primarily via *MONOPTEROS (MP/ARF5), ARF7* and *ARF19*^4^.We therefore treated two different allelic *arf7,19* mutant combinations and the conditional triple mutant *amiMP* (inducible artificial microRNA against *MP* in *arf7,19*^4^; see Methods) with GA. No significant changes in the number of secondary xylem vessels were observed in any of the mutant combinations following GA treatment (**Fig. 3e,f**), indicating that GA’s effect on xylem production requires *ARF5/ARF7/ARF19-mediated* auxin signalling.

The HOMEODOMAIN LEUCINE ZIPPER IIIs (HD-ZIP IIIs) act downstream of auxin signalling^31,32^ to promote xylem identity in the root cambium^4^. A representative member of the family, *AtHB8*, is expressed specifically in the early xylem cells^4^. Since *ga1* has a narrow auxin signalling maximum (**Fig. 3b; Extended Data Fig 2a**), *AtHB8* expression is also reduced in the *ga1* mutant, as shown by qRT-PCR analysis (**Fig. 3d**). Inducible overexpression of *mir165*, which targets the mRNAs of all five HD-ZIP IIIs for degradation^33^, leads to the inhibition of secondary xylem formation in the root cambium^4^. GA was unable to rescue this phenotype, indicating that the HD-ZIP IIIs are required for GA-induced xylem production (**Fig. 3g,h**). Taken together, these data show that GA’s effect on xylem formation acts via auxin signalling and its downstream factors to define xylem identity.

### GA promotes long distance PAT via PIN1

The PIN auxin efflux carriers play a dominant role in determining how auxin accumulates in different tissues^34^. Since GA has previously been reported to regulate PIN levels in the root apical meristem^22,35^, we investigated whether GA also regulates auxin accumulation, and thus auxin signalling, through PIN activity in the vascular cambium. Of the five plasma membrane-localized PINs (PIN1,2,3,4,7)^34^, only PIN1 showed consistent expression on the xylem side of the vascular cambium **(Extended Data Fig. 3a-e)**.A detailed analysis revealed that PIN1 has the highest expression in the xylem-side stem cell daughters (position −1), with weaker expression in the neighbouring cells (positions −2 and 1). Following 24 h GA treatment, PIN1 expression spreads towards the phloem to occupy both stem cell daughters (**Fig. 4a,c**), thus showing a shift in expression similar to the auxin signalling marker *DR5v2*. However, *DR5v2* induction takes longer time (48 h) (**Fig. 3a-c; Extended data Fig. 2b-d**). In the *ga1* mutant background, the PIN1 expression pattern is similar to the pattern in wild type, but a similar shift in PIN1 expression was observed after GA treatment (**Fig. 4b,c**). Together, these data show that GA promotes PIN1 expression in the stem cells and this is followed by expression of *DR5v2*.

**Figure 4.**
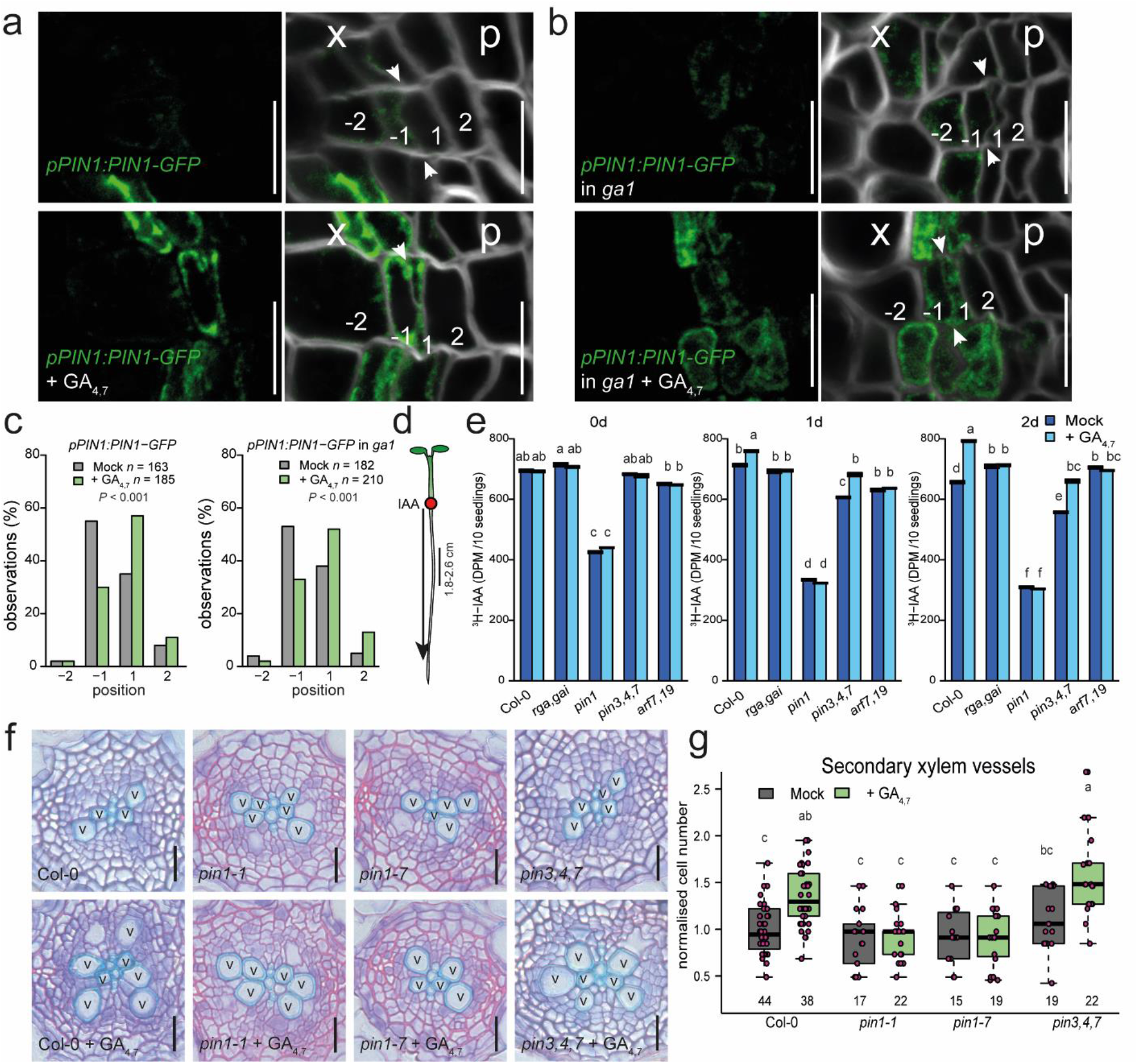
GA promotes long-distance auxin transport in a PIN1-dependent manner. Expression of *pPIN1:PIN1-GFP* in the root cambium after a 24 h GA treatment in 14-day old seedlings of wild type (**a**) and *ga1* (**b**). White arrowheads indicate the most recent cell divisions. The numbers “-2”, “-1”, “1” and “2” indicate the position of the cells relative to the most recent cell division, with negative values towards the xylem and positive towards the phloem. (**c**) Counts of the position in the cambium at which the *pPIN1:PIN1-GFP* gradient ends. Cellular positions on the x-axis correspond with the cellular positions in panels **a** & **b**, and n refers to the total number of observations. (**d**) A schematic explaining the setup of the PAT assay. The red circle indicates the position of ^3^H-IAA application, black arrow showing the direction of IAA movement. The black line indicates the area sampled to detect ^3^H-IAA. (**e**) ^3^H-IAA transport from the root-shoot transition zone to 1.8-2.6 cm from the root tip after a 1 h GA treatment in 6-day old Col-0 and mutant plants. After 0, 1, or 2 days, plants were treated with ^3^H-IAA for 3 h and then sampled. Data shown are means ± SD (n = 3 independent pools of 10). (**f**) Root cross-sections after a 6-day GA treatment in 4-day old seedlings of Col-0 and various *pin*-mutants. (**g**) Quantification of the number of secondary xylem vessels in plants shown in panel **f**. Chi-squared test in **c**; two-way ANOVA with Tukey’s post hoc test in **e** and **g**. The boxes in the box and whisker plots show the median and the interquartile range, and the whiskers show the total range. Individual data points are plotted as purple dots. Numbers in **g** indicate number of samples. Letters indicate a significant difference, *P* < 0.05. Scale bars are 10 μm (**a,b**) or 20 μm (**f**). “p”= phloem, “x”= xylem, “v”= secondary xylem vessel. All experiments were repeated three times.

Previously, PIN1 has been proposed to act both via increased long distance PAT and via local redirection of auxin fluxes^11,13,34^. PIN1 has been shown to be basally localised in vascular cells^11,13,36,37^. Similarly, in the root cambial stem cells, we observed basal PIN1 localisation, which did not change after GA treatment (**Extended Data Fig. 3f**). This suggests that GA does not redirect auxin fluxes within the cambium, implying that long distance PAT might be affected. To test whether GA enhances long distance PAT, we performed a PAT assay. 6-day old seedlings were treated with GA4,7 for 1 h, after which seedlings were rinsed and then transferred either directly to discontinuous media for auxin transport assay or replaced on MS media to grow for an extra one or two days. For the PAT assay, tritium labelled indole-3-acetic acid (^3^H-IAA) was applied to the root-shoot transition zone, and radioactivity was measured in either the upper part of the root (**Fig. 4d,e**) or the root tip (**Extended Data Fig. 4a,b**). Increased ^3^H-IAA signals were observed in the upper part of GA-treated wild type roots one day after GA application (**Fig. 4e**). As expected, in the DELLA double mutant *rga,gai*, in which GA signalling is derepressed^38^, the ^3^H-IAA signal did not increase upon GA treatment (**Fig. 4e; Extended Data Fig. 4b**), thus demonstrating that GA’s effect on PAT is caused by the canonical GA signalling pathway. Similarly, *arf7,19* failed to respond to GA (**Fig. 4e; Extended Data Fig. 4b**), indicating that ARF7/19-mediated auxin signalling is required for GA-induced PAT as well as for xylem formation (**Fig. 3e,f**).

As GA signalling is able to both enhance PAT and broaden PIN1 expression in the cambium, we postulated that PIN1 might be required for GA’s effect on PAT. The *pin1-7* loss-of-function mutant has a lower baseline level of PAT, and *pin1* mutant roots did not show increased ^3^H-IAA transport upon GA treatment, similar to *rga,gai* and *arf7,19* mutants (**Fig. 4e** and **Extended Data Fig. 4b**). However, GA treatment in the triple mutant lacking three of the other plasma membrane localised PINs, *pin3,4,7*, did result in increased levels of ^3^H-IAA in roots (**Fig. 4e** and **Extended Data Fig. 4b**), indicating that mainly PIN1 is required for GA’s effect on long-distance PAT.

In addition to PINs, two ATP Binding Cassette subfamily B (ABCB) auxin transporters, ABCB19 and ABCB21, also contribute to maintenance of polar auxin transport streams in the vasculature^39,40^. No change in *ABCB19* expression was observed with GA treatment (**Extended Data Fig. 5a**). However, *ABCB21*, which is localised almost exclusively to the pericycle^40^, initially increased slightly with GA treatment and maintained over a 24 h period (**Extended Data Fig. 5a,b**). While rootward auxin transport was severely reduced in *abcb19*, mutants still showed increased transport with GA treatment (**Extended Data Fig. 5c**). PAT in *abcb21* was only slightly responsive to GA (**Extended Data Fig. 5c**). Together these results suggest that GA-enhanced long-distance PAT requires ABCB19 function along with PIN1. Additionally, GA-upregulated ABCB21 likely increases restriction of auxin to the central vasculature, where PIN1 provides directional flux toward the root tip in addition to more localized auxin distributions within vascular cambium.

Since PIN1 has a central role in directional auxin flux along cambium, we next studied whether PIN1 is required for GA to promote secondary xylem production. We first analysed the effect of GA treatment in two allelic *pin1* mutants, *pin1-1* and *pin1-7*. GA treatment led to an increased number of secondary xylem vessels in wild type but not in either of the *pin1* mutants (**Fig. 4f,g**). In contrast, the *pin3,4,7* mutant responded similarly to wild type in terms of xylem production (**Fig. 4f,g**), indicating a non-redundant function for PIN1 in GA-induced xylem formation. Altogether, our data show that GA promotes broadening of PIN1 expression in the cambium, which results in increased PAT along the hypocotyl and root. This leads to a broadening of the high auxin signalling domain in cambium, thus promoting xylem production.

### GA treatment occasionally leads to stem cell respecification

Next, we investigated how the GA-induced changes in the width of the auxin maximum alter stem cell fate decisions, shifting from equal xylem and phloem distribution towards favouring xylem production (**Fig. 1f-i**). First, we investigated the stem cell division dynamics using the stem cell marker *pANT:erRFP* together with labelling dividing cells with 5-ethynyl-2’-deoxyuridine (EdU)^41^. *ANT* was typically expressed in both stem cell daughters (mock: 68%; **Fig. 5a**) and to a lesser degree only in the phloem-side stem cell daughter (32%). After two days of EdU tracing, the majority of the EdU-positive cells were in the *ANT* expression domain (mock: 80%; **Fig. 5b**). However, following a 2-day GA treatment, a larger proportion of *ANT* expression was restricted to the phloem-side stem cell daughter (GA4,7: 43%; **Fig. 5a**). In addition, significantly more EdU-positive cells were outside the *ANT* expression domain towards the xylem (mock: 20%, GA: 36%; **Fig. 5b**). These data show that GA treatment results in a higher proportion of xylem-side stem cell daughters losing stem cell identity and obtaining xylem identity.

**Figure 5.**
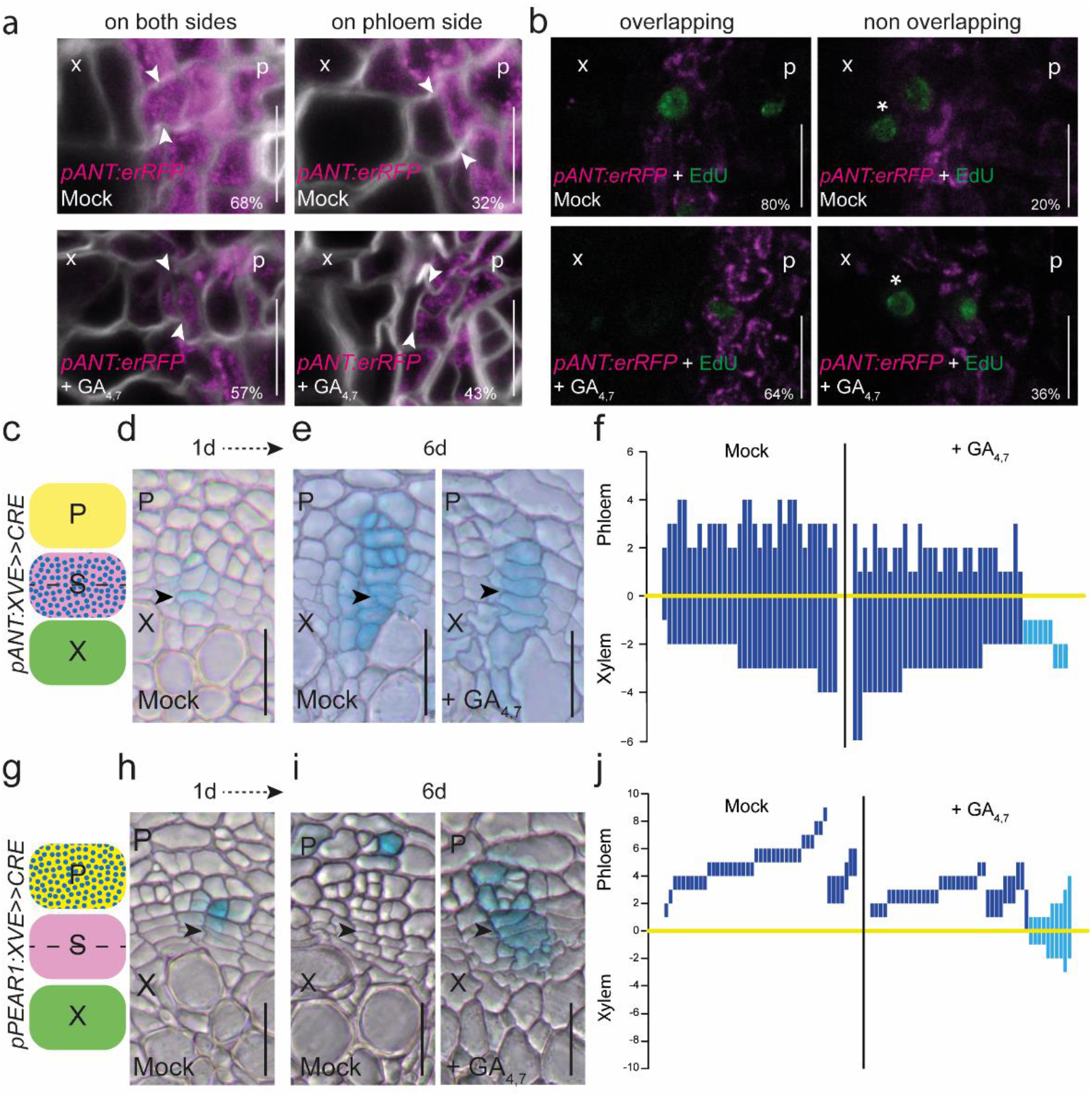
GA promotes xylem formation by influencing cambial dynamics. (**a**). Confocal root cross-sections of *pANT:erRFP* after a 48 h GA treatment in 14-day old seedlings. White arrowheads indicate the most recent cell divisions. A shift of expression to only the phloem-side stem cell daughter after GA treatment was significant, with a *P* value of 0.0012 (chi-square test, nmock=407, nGA= 477). (**b**) Confocal root cross-sections of *pANT:erRFP* (magenta) after 6 hours of EdU (green) incorporation and a 48 h GA treatment in 14-day old seedlings. White asterisks indicate EdU-positive cells that do not overlap with *pANT:erRFP* expression. The increase in EdU-positive nuclei not overlapping with *ANT* expression was significant, with a *P* value of 0.03 (chi-square test, nmock=100, nGA= 102).The percentages in the corners of the subpanels represent the frequency of the observed phenotypes (**a, b**). (**c**) A schematic showing where *ANT* sectors originate from within the vascular cambium. (**d**) An example of a stem cell sector one day after induction in a 14-day old seeling. (**e**) Examples of stem cell sectors 6 days after induction in 14-day old seedlings grown on 2 μM GA4,7 or mock treatment. (**f**) GUS sectors (bars) plotted based on the position of the thinnest cell wall (yellow line). Note that the light blue bars highlight the sectors that are only present on the xylem side of the cambium (21% of the GA treated samples). (**g**) A schematic describing where the *PEAR1* sectors originate from within the vascular cambium. (**h**) An example of a phloem cell sector one day after induction in a 14-day old plant. (**i**) Examples of the phloem sectors 6 days after induction in 14-day old plants grown on 2 μM GA4,7 or mock treatment. (**j**) GUS sectors (bars) plotted based on the position of the thinnest cell wall (yellow line). Note that the light blue bars highlight the sectors that are able to produce both xylem and phloem (21% of the GA treated samples). “x”= xylem, “p”= phloem, “S”= stem cell. Black arrowheads indicate the most recent cell divisions. Percentages in **a** and **b**indicate frequency of the observed phenotype. Scale bars are 20 μm (**d**, **e, h, i**) or 10 μm (**a**, **b**). All experiments were repeated three times.

In order to follow the consequences of altered stem cell dynamics during long-term GA treatment, we carried out a lineage tracing experiment where sectors marked with GUS expression were induced in the stem cells using the *ANT* promoter (**Fig. 5c,d**)^4^. Under normal growth conditions, stem cell sectors spanned almost equally towards both the xylem and the phloem (**Fig. 5e,f**), similar to the stem cell sectors generated randomly within the cambium (**Fig. 1f,g,i**) and what we have shown earlier^4^. When seedlings are treated with GA, the majority of the stem cell sectors spanned further towards xylem than phloem (**Fig. 5e,f**). Unexpectedly, a subset of the ANT-sectors were pushed away from the cambium into the xylem (light blue sectors in **Fig. 5f**, 21% of the GA sectors), indicating that, occasionally, both stem cell daughters lose their identity and differentiate into xylem after GA application. This led us to hypothesise that when auxin signalling spreads to both stem cell daughters causing them to differentiate into xylem, the adjacent phloem identity cell respecifies as a stem cell. To test this, we performed a lineage tracing experiment with sectors originating from a single early phloem cell using the promoter of phloem identity gene *PEAR1*^29^ (**Fig. 5g,h**). Under normal conditions, the active cambium pushes phloem identity cells away from the cambium while they differentiate into phloem cells, leading to the formation of sectors deep in the phloem (**Fig. 5i,j**). However, under GA-treated conditions, a subset of phloem lineage sectors is able to produce both xylem and phloem (light blue sectors in **Fig. 5j:**, 21% of the GA sectors), indicating that in these sectors the lineage progenitor re-acquired stem cell identity. These data suggest that the original phloem identity cell occasionally respecifies as a stem cell during GA treatment, thus supporting the respecification hypothesis.

## Discussion

We show that GA affects xylem proliferation in two ways: first, it increases the number of xylem cells differentiating from the stem cells, and second, it promotes the expansion of secondary xylem vessels, resembling the effect that GA has on other cell types in other tissues^42^. GA has the opposite effect on phloem production: stem cells produce fewer phloem cells. However, despite the reduced total phloem cell number, a higher proportion of conductive cells are produced. In turn, a GA biosynthesis mutant has a higher proportion of parenchyma cells than conductive cells. Thus, even though GA levels have a clear impact on phloem production, they have a smaller impact on the number of conductive phloem cells. This might be important in ensuring phloem transport capacity regardless of GA status. Auxin promotes primary sieve element differentiation in root tips^43^. Since we show that GA increases auxin signalling in cambium and that GA also promotes conductive phloem formation, we speculate that auxin is needed for the differentiation of conductive phloem cell types also during secondary growth. Supporting this hypothesis, studies have shown that GA and auxin together increase the production of phloem fibres^44,45^.

We discovered that GA promotes PIN1-dependent and ABCB19/21-assisted PAT, which leads to elevated auxin accumulation and signalling in the root cambium during the early stages of secondary development. Previous studies have shown that the DELLAs and ARFs together regulate xylem production in the *Arabidopsis* hypocotyl during flowering^21^. In poplar stems, GA promotes xylem production via *ARF7*, and this is associated with transcriptional upregulation of *PIN1*^46,47^. During leaf venation, PIN1 promotes auxin accumulation^48^, which leads to activation of ARFs^49^. This in turn promotes *PIN1* expression, thus completing a feed-forward loop^50^. Our results show that GA induces PIN1 first, followed by upregulation of the ARF-regulated auxin signalling reporter *DR5v2*. These results support a mechanism in which GA enters this feed-forward loop by regulating the PIN1 expression pattern, at least during early secondary development in the *Arabidopsis* root.

Organizer cells in meristems position the stem cells to the adjacent cells. In the cambium, organizer cells are defined by a local auxin signalling maximum and subsequent HD ZIP III expression that leads to cells acquiring xylem identity and cell-autonomous inhibition of cell division^4^. In this study, we show that the position of the maximum regulates the fate decisions of the stem cell daughters. In the presence of high GA and thus elevated PAT, the xylem-side stem cell daughters accumulate high levels of auxin and therefore likely obtain xylem/organizer identity. The phloem-side stem cell daughters retain stem cell identity (**Fig. 6a**). Occasionally, both daughters accumulate high levels of auxin, leading both to obtain xylem/organizer identity. This forces the neighbouring phloem identity cell to respecify as a stem cell (**Fig. 6b**). When GA levels are low, both stem cell daughters have low auxin levels, thus making the xylem-side daughter maintain its stem cell identity, since it is located adjacent to an existing auxin signalling maximum. Under these conditions, the phloem-side daughter obtains phloem identity. It is unknown what positions the stem cells adjacent to the auxin maximum. One possibility is that medium auxin levels within the auxin gradient promote stem cell divisions. Supporting this idea, we previously observed an auxin signalling gradient along the cambium using a sensitive auxin signalling reporter^4^. However, it is unclear how such a gradient could robustly position the stem cells. Another possibility is that the auxin maximum initiates a mobile signal, which non-cell-autonomously specifies stem cells in the adjacent position and promotes their division. However, the existence of such a signal remains speculative.

**Figure 6.**
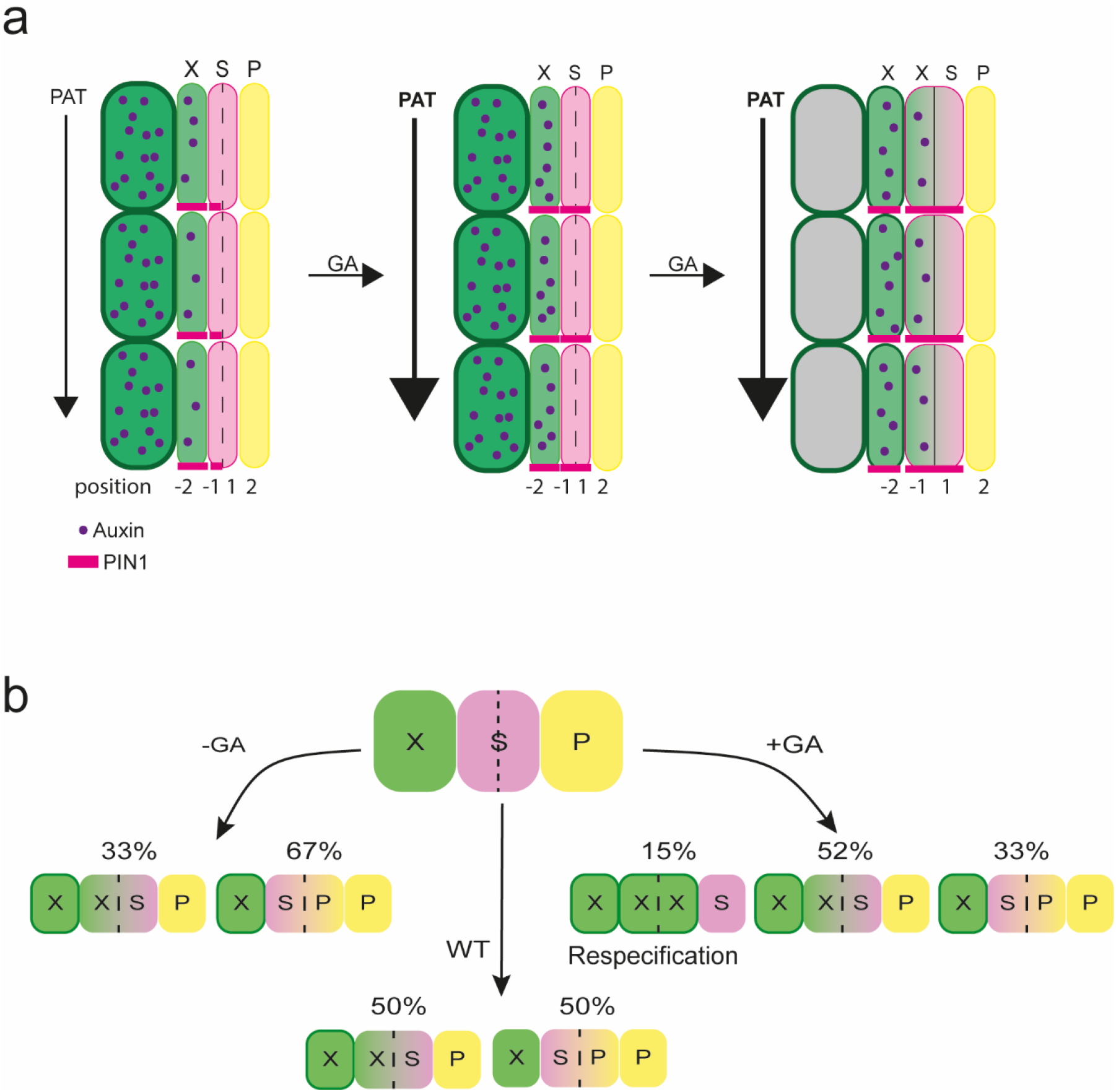
Models describing cambium dynamics. (**a**) Model showing what happens to PAT, PIN1 and auxin signalling upon GA treatment. Increased PAT induces PIN1 in the phloem-side stem cell daughter, and this leads to the widening of the auxin signalling maximum to the xylem-side stem cell daughter, which then gains xylem identity. The numbers “-2”, “-1”, “1” and “2” indicate the position of the cells relative to the most recent cell division, with negative values towards the xylem and positive towards the phloem. (**b**) Model explaining how the fate of the stem cell daughters is regulated by GA. In normal conditions, cambial stem cells produce an equal amount of xylem and phloem. With low GA levels, stem cell daughters preferentially gain phloem identity, while high GA levels lead to xylem identity, and in extreme cases the respecification of stem cells from phloem identity cells. X = xylem, S = stem cell (daughters), P = phloem.

## Acknowledgements

We would like to thank Claus Schwechheimer, Enrico Scarpella and Laura Ragni for providing us published material; and Laura Ragni, Enrico Scarpella, Sedeer el-Showk, Hiroyuki Iida and Xixi Zhang for providing feedback on the manuscript. Confocal imaging was performed with help and using equipment of the Light microscopy Unit (LMU), University of Helsinki. Special thanks to Mikko Herpola and Miki Iida for helping with daily lab related tasks. This work was supported by the Academy of Finland (grants #316544, #346141), European Research Council (ERC-CoG CORKtheCAMBIA, agreement 819422), University of Helsinki (HiLIFE fellowship and DPPS) and the US Department of Energy, Basic Energy Sciences, grant no. DE-FG02-06ER15804 to A.S.M. and M.K.J.

## Contributions

A.P.M. conceived the project; A.P.M., R.M. and O.S. designed the experiments; R.M., O.S. and B.W. performed the experiments, except M.K.J and A.S.M. designed and conducted the PAT experiment and analysis of ABCBs; L.V. created the APL projections; A.S.G. provided preliminary data; M.L., L.Y., X.W. and R.S. generated genetic material; A.P.M, R.M. and B.W. wrote the paper with input from all authors.

## Data availability

All data supporting the findings of this article are available in this article and its supplementary information. Source data are provided with this paper.

## Methods

### Gene accession numbers

The accession numbers of the genes in this study are: *CYCB1;1*, AT4G37490; *PEAR1*, AT2G37590; *ANT*,AT4G37750; *AtHB8*, AT4G32880; *MIR165A*, AT1G01183; *MP*, AT1G19850; *ARF7*, AT5G20730; *ARF19*,AT1G19220; *GA1*, AT4G02780; *RGA*, AT2G01570; *PIN1*, AT1G73590; GAI, AT1G14920; *PIN2*, AT5G57090; *PIN3*, AT1G70940; *PIN4*, AT2G01420; *PIN7*, AT1G23080; *APL*, AT1G79430; ABCB19, AT3G28860; ABCB21, AT3G62150.

### Plant material and cloning

All entry clones, except p1R4z-DR5v2, were generated by PCR amplification of the desired sequence with the primers listed in Table 1 followed by recombination into Multisite Gateway compatible pDONR entry vectors (Table 2). The PCR fragment of *DR5v2*, which was amplified from genomic DNA isolated from *DR5v2:nlsGFP*^30^,was cloned into the *p1R4z-BsaI-ccdB-BsaI* entry vector via Golden Gate cloning to generate *p1R4z-DR5v2*.The construction of *p1R4z-BsaI-ccdB-BsaI* and the Golden Gate cloning were done as previously described^51^. The resulting entry vector, *p1R4z-DR5v2* was assembled together with *p221z-erRFP*^52^ and *p2R3z-nosT*^52^ into the destination vector *pHm43GW*^53^ by a MultiSite Gateway LR reaction.

Multisite Gateway technology was used to combine entry clones carrying a promoter (1st box), gene of interest or a tag (2nd box) and a tag or terminator (3rd box) with Gateway-compatible binary destination vectors in a multisite Gateway LR clonase reaction. All of the expression vectors generated in this study are listed in Table 3.

All of the expression vectors were dipped in the Col-0 background, and single insertion lines were screened based on Mendelian segregation of the selection marker. Several single insertion lines were screened for each construct to observe the most consistent phenotypes or expression patterns. A previously published inducible miRNA against MP (*amiMP*)^4^ line was dipped into the *arf7-2,19-5* background due to silencing issues in the earlier *arf7-1,19-1* background. Seeds published in this study, as well as the already published lines, are listed in Supplementary Table 4. The following transgenic and mutant lines have been reported elsewhere: *pHS:Dbox-CRE x 35S:lox-GUS^4^, p35S:XVE>>miR165a^4^, pANT:XVE-CRE x 35S:lox-GUS^4^, pPIN1:PIN1-GFP^54^, pPIN1:PIN1-GFP x ga1^22^, pPIN2:PIN2-GFP^54^, pPIN3:PIN3-GFP^55^, pPIN4:PIN4-GFP^56^, pPIN7:PIN7-GFP^56^, pRGA:GFP-RGA^27^, arf7-1,19-1^57^, arf7-2,19-5^58^, pin1-7 (SALK-047613)^59^, pin3,4,7^60^, pin1-1^11^, ga1 (SALK-109115*)^22^, *abcb19-101*^61^ and *abcb21-1*^40^.

### Plant growth and chemical treatments

Seeds were surface sterilised first with 20% chlorine and then with 70% ethanol, washed twice with H2O and then plated on a half-strength growth medium (½ GM, containing 0.5 × MS salt mixture with vitamins (Duchefa), 1% sucrose, 0,5g/l MES pH 5.8 and 0.8% agar) and vernalized at 4 °C for 2 days. In the case of *ga1* (SALK-109115), after sterilisation the seeds were soaked in 100 μM GA3 for 5 days and covered at 4 °C. Before plating, seeds were washed 5 times with H_2_O. The age of the plants was measured from when the plates were vertically positioned in the growth cabinet. The temperature in the cabinets was 22 °C and they had long-day conditions (16h of light). In order to get seeds from *ga1* plants, plants growing in soil were sprayed with 100 μM GA3 twice per week until they had seeds.

10 mM and 100 mM stocks of GA_4,7_ (Duchefa) and GA3 (Duchefa) were prepared in 100% EtoH and stored at −20°C. A 10 mM stock of EdU, a thymidine analogue (Thermo Fisher), was made in DMSO and stored at −20 °C. 17-b-oestradiol (Sigma), a synthetic derivative of oestradiol, was prepared as a 20 mM stock solution in DMSO and stored at −20 °C.

100 μM GA_3_ was used for *ga1* seed germination and seed production. The working concentration for GA_4,7_ was 2 μM. XVE-based gene induction was achieved by transferring plants onto plates containing 5 μM 17-b-oestradiol or an equal volume of DMSO as a mock treatment. For EdU incorporation, plants were placed in liquid ½GM containing 10 μM EdU for the time stated in each experiment.

### GUS-staining, microtome sections and histology

The protocol was modified from Idänheimo et al.^62^. Samples were fixed with 90% acetone on ice for 30 min, washed two times with a sodium phosphate buffer (0.05 M, pH 7.2) and then vacuum infiltrated with the GUS-staining solution (0.05 M sodium phosphate buffer, pH 7.2; 1.5 mM ferrocyanide, 1.5 mM ferricyanide, 1 mM X-glucuronic acid, 0.1% Triton X-100). Samples were placed at 37 °C until the staining was at the desired level (the required time varied between different lines).

After staining, the samples were fixed overnight in 1% glutaraldehyde, 4% formaldehyde, and 0.05 M sodium phosphate pH 7.2. Fixed samples were dehydrated in an ethanol series (10%, 30%, 50%, 70%, 96%, 2x 100%), with 30 minutes for each step, and then incubated for 1 h in a 1:1 solution of 100% ethanol and solution A (Leica Historesin Embedding kit). After 2 h in solution A, samples were placed in plastic chambers and filled with 14:1 mixture of solution A: hardener.

Sections of 5 or 10 μm were prepared on a Leica JUNG RM2055 microtome using a microtome knife (Leica Disposable blades TC-65). The sections were imaged without staining or after staining with Safranin O (Sigma-Aldrich) (1 min in 0,0125% solution, rinsed with water) or double staining with 0.05% Ruthenium Red (Sigma–Aldrich) and Toluidine blue (Sigma Aldrich) (5 s in each, rinsed between stainings and afterwards with water). Sections were mounted in water and visualised with a Leica 2500 Microscope.

### Fluorescent marker analysis: vibratome sections and EdU detection

Using a protocol modified from Smetana et al.^4^, samples were vacuum infiltrated with 4% paraformaldehyde solution (PFA, Sigma) in 1xPBS pH 7.2. After fixation, the samples were washed with PBS and embedded in 4% agarose. Embedded samples were cut with a vibratome into 200 μm sections for confocal analysis. Agarose slices were placed into PBS with SR2200 (1:1000, Renaissance Chemicals) for cell wall staining. For root tip visualizations, we fixed the samples with 4% PFA, cleared them with CLEARSEE, and stained the cell walls with SR2200 as in Ursache et al.^63^.

To visualise EdU-positive nuclei, EdU detection was performed on the agarose sections before cell wall staining. The Click-iT EdU Alexa Fluor 488 Imaging Kit (Thermo Fisher) was used for detection with a modified EdU detection mix^41^. Samples were incubated in the detection mix for 1 h on ice and then transferred into PBS with SR2200 (1:1000).

### Microscopy and image processing

Light microscopy images were taken with a Leica 2500 microscope (20x and 40x objectives). Fluorescent markers were imaged with a Leica Stellaris 8 confocal microscope. Confocal images were obtained with Leica Las AF software using PBS or water as the imaging medium. All confocal images with multiple channels were imaged in sequential scan mode. Confocal settings may have varied between experiments but always stayed the same for the experimental sample and the respective control. In order to better optimise the SR2200 cell wall staining, the signal was sometimes adjusted during imaging and may thus vary between the sample and control.

The Leica Stellaris 8 has a Tau-gating mode that makes it possible to separate GFP signals from background signals. GFP markers were always imaged with this Tau-gating mode, gathering signals between 1.3-9 ns.

### Image projections

For image projections (**Extended Data Fig. 1c-e**), each image was annotated by marking the centre of the root and following the most recent cell division in each cell column in the cambium. The images have been rotated so that the primary xylem axis is oriented in vertical position. Signal data from the image was sampled from the centre point to the edges of the root and aligned to the most recent cell division in the cambial zone. All images in the same treatment were then aligned with the annotated cambial line starting from the centre to the edge. Images within each treatment can therefore be compared and analysed based on the fluorescent signal distribution and intensity and the location/distance of cambium from the root centre. Image wrapping was done using Python 3.8.10^64^ and image ROI area extraction was done using several different libraries, including OpenCV2^65^, Pillow^66^, Matplotlib v2.2.1^67^ and NumPy^68^. More detailed documentation is available on Github (https://github.com/LMIVainio/PolarUnwrap/find/main).

### Image analysis

Fiji/ImageJ was used for image analysis. When counting secondary xylem vessels, the primary xylem axis was not included and only mature secondary vessels with light blue toluidine blue staining were counted. Cells were counted with the cell counter tool. Xylem cells and include all the cells inwards of the most recent (=thinnest) cell division, so this also sometimes includes the stem cells and stem cell daughters (black line in **Fig. 1A**). Phloem cells were counted as all the cells outwards from the most recent cell division until the periderm border (clearly thicker continuous cell wall on the outskirts of the cross section: red line in **Fig. 1A**). In **Fig. 4f,g** (pin mutants), the data in the graph is combined from 4 separate experiments, so we normalised the data from the experiments by giving the control (Col-0) the value of 1 and counting the other values relative to that.

Analysis of the fluorescent markers was done with either Fiji/ImageJ or Leica LAS X lite. For *PIN1* and *DR5*, we quantified the reach of the respective marker expression, meaning the position of the last cell in cambium marker expression was seen. For the spread of *ANT*, we quantified the expression of the *ANT* marker in the cambium, recording whether the marker was expressed on both sides of the most recent cell division or only on the phloem side. Both of these quantifications were only done on cell lineages where the thinnest cell wall was clearly recognisable. For the EdU pulse experiment, we quantified the number of EdU positive nuclei that either overlapped with the *ANT* signal or were on its xylem side, and the number of those which are only on the xylem side of *ANT* expression.

### Auxin transport assays

6-day old seedlings on ½ MS agar plates were treated by applying a thin surface drench of 3 μM GA_4,7_. After 1 hour, the solution was poured off and the seedlings were rinsed and gently blotted to remove excess solution. The seedlings were then either transferred directly to a discontinuous filter paper system for transport assays^69–71^ or allowed to grow for an additional 1-2 days prior to the assays. For the auxin transport assays, a 200 nL droplet of 10 μM ^3^H-IAA was placed at the root-shoot transition zone and the seedlings were then incubated under low yellow light. After 3 hours, 8 mm segments were collected from two different positions along the root: apex-0.8 cm (=root tip) and 1.8-2.6 cm (=upper part). ^3^H-IAA was measured by liquid scintillation counting. The 1.8-2.6 cm segments contained lateral root primordia and emerged lateral roots. Data shown are means ± SD (3 independent pools of 10 seedlings).

### qRT-PCR

RNA was collected from 2 cm long pieces starting just below the hypocotyl of 10-day old roots where lateral roots had been removed. RNA was isolated using the GeneJET Plant RNA Purification Mini kit (Thermo Fisher) and treated with DNAse. cDNA was synthesised from 100 ng of RNA using Maxima H Minus reverse transcriptase (Thermo Fisher) and oligodT primers (Thermo Fisher). The PCR reaction was done on a Bio-Rad CFX384 cycler using EvaGreen qPCR mix (Solis Biodyne) and the following program: 10 min at 95 °C, 50 cycles (10 s at 95 °C, 10 s at 60 °C, 30 s in 72 °C). All of the primers used in qRT-PCR are listed in Table 1. The results were normalised, following earlier published methods^72,73^, to the reference genes *ACT2, UBQ10* and *TIP41*. Three biological replicates were used for each line and treatment, as well as three technical replicates.

For ABCB21 expression, 7d seedlings were surface drenched with MS solution containing solvent control, 1 μM, or 10 μM GA for 15 mins. Solutions were decanted then plates returned upright in light for 24h. Total RNA was isolated with TRIzol (Thermo Fisher) followed by lithium chloride precipitation. 1.5 μg total RNA was reverse transcribed with Superscript III (Thermo Fisher). PCR reactions were performed on a Bio-Rad CFX96 cycler using SYBR Green master mix (Applied Biosystems) and the following program: 3 m at 95°C, 45 cycles (15s at 95°C, 1 min at 60°C). Expression was normalized to the reference genes ACT2 and PP2A. Primers used were from Jenness et al., (2019)^40^.

### ANT EdU pulse experiment

A short 6 h pulse of 10 μM EdU in liquid ½GM was used, after which the EdU was removed by washing twice for 15 min with liquid ½GM. Washed plants were transferred into 2 μM GA4,7 or EtOH plates and allowed to grow for 2 days. After this, they were fixed for agarose sections and confocal analysis.

### Lineage tracing

All lineage tracing experiments were performed in 16-day old plants. For the *pHSdboxCRE* plants, plates were placed at 37°C for 14 or 17 minutes. They were then immediately cooled at 4 °C for 15 minutes^4^. The plants were then transferred to 2 μM GA4,7 or EtOH plates for 6 days. For the oestradiol-inducible lineage tracing lines, plants were incubated in 5 μM EST in liquid ½GM for two hours (*pPEAR1:XVE>>CRE*) or 30 min (*pANT:XVE>>CRE*), washed 3x 15 min and then transferred to 2 μM GA4,7 or EtOH plates for 6 days. For the *pHSdboxCRE* experiments, we considered for the analysis only the sectors that proliferated.

### General methodology and statistical analysis

The number of individual plants, cross sections or clones analysed is displayed as the n in figures or figure legends. The fraction in the corner of some images indicates the frequency of the observation. All statistical analyses were performed using R version 4.1.2 (http://www.r-project.org/).

All measurements were taken from distinct samples and the same sample was not measured repeatedly.

Before comparing means, the normality of the data was confirmed with the Shapiro-Wilk test. When doing pairwise comparisons, normally distributed data were analysed with a 2-tailed t-test and other data with a 2-tailed nonparametric Wilcoxon test. When comparing multiple means with each other, a two-way ANOVA with Tukey post hoc was performed. Categorical data were analysed with a chi-squared test.

In all of the box plots, the centre line represents the median, and the upper and lower box limits indicate the 75th and 25th percentiles, respectively. Whiskers show the maximum and minimum values, and outliers are shown as circles. Filled circles represent individual data points. In violin plots, the white dot shows the median and the thick line the interquartile range. The thinner line represents the rest of the distribution. Each side of the line is a kernel density estimation that shows the distribution shape of the data. Filled circles represent individual data points.

### Softwares used

Leica LAS x, Leica LAS x lite, Bio-Rad CFX Manager, Fiji 1.53, R 4.1.2, R-studio, Adobe Illustrator, Python 3.8.10, MS Office: Excel, Word

## Extended Data Figures

**Extended Data Figure 1.**
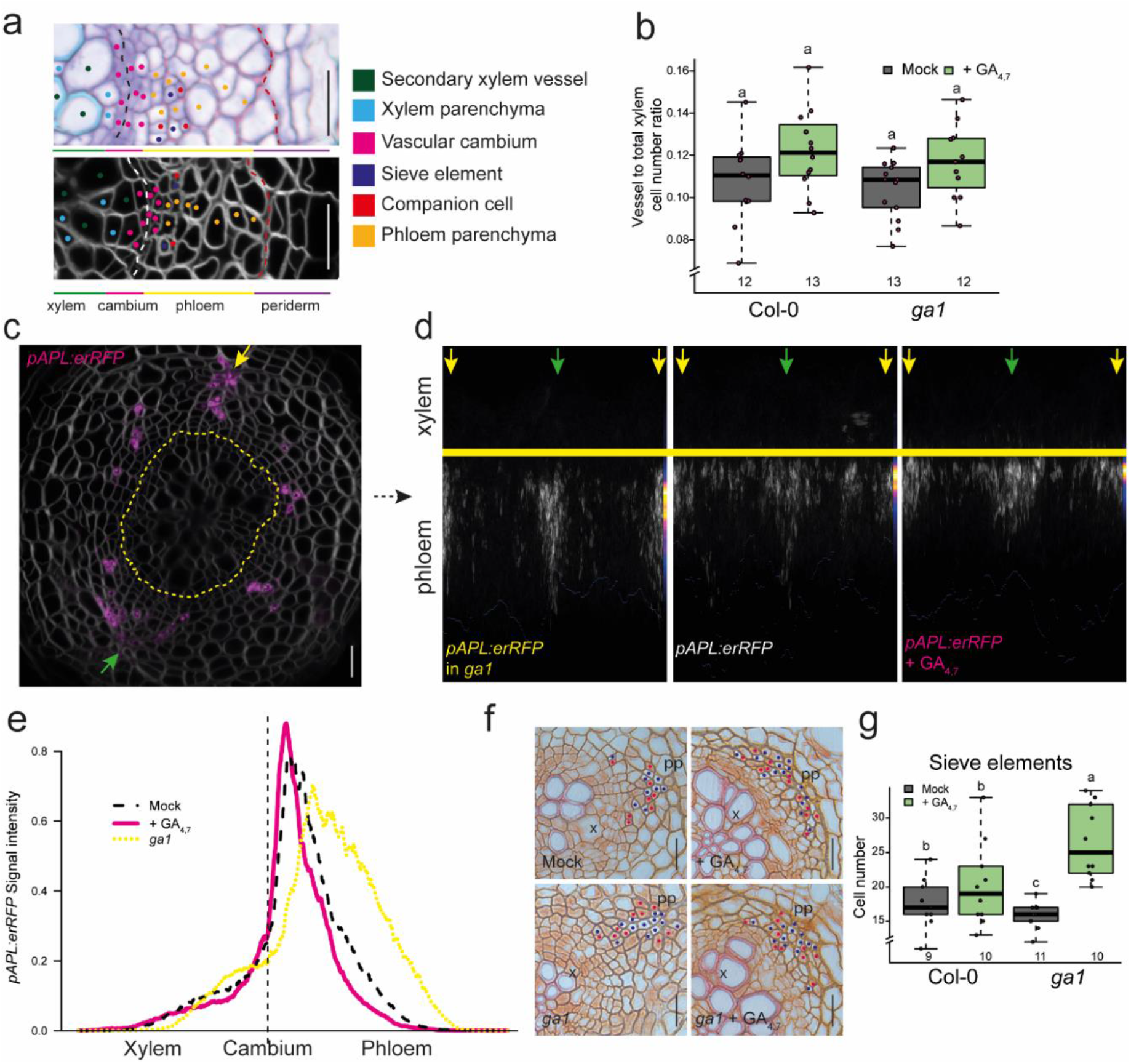
Characterisation of secondary tissues after GA treatment. (**a**) Schematic describing secondary growth tissue and cell types in plastic and agarose sections of 14-day old roots. Black dotted lines indicate the most recent cell divisions. Red dotted lines mark the border between the phloem parenchyma cells and the periderm. (**b**) The ratio of secondary xylem vessels to total xylem cell number. (**c**) An example of *pAPL:erRFP* expression in 14-day old roots. The dashed yellow line marks the most recent cell divisions. (**d**) Projections of *pAPL:erRFP* roots of 4-day old plants grown for 10 days on mock or 2 μM GA_4,7_ plates or crossed into *ga1*. Each picture is combined from ~15 pictures with the phloem poles and thinnest cell walls aligned. The cambium is marked by a yellow line. Yellow and green arrows point to the primary phloem poles. Heat maps on the side show where the expression accumulates. (**e**) Graph showing *APL* expression relative to the cambium position. (**f**) Safranin-stained cross-sections of Col-0 and *ga1* 4-day old plants treated for 10 days with 2 μM GA4,7 or mock. Safranin O does not stain sieve elements (blue dots), thus they stay white and are easy to distinguish. Companion cells are marked with red dots. (**g**) Quantification of the number of sieve elements in safranin-stained cross-sections. “x” = xylem, “pp”= primary phloem pole. Two-way ANOVA with Tukey’s post hoc test in **b, g**. The boxes in the box and whisker plots show the median and interquartile range, and the whiskers show the total range. Individual data points are plotted as purple dots. Letters indicate significant differences, with *P* < 0.05. Scale bars are and 20μm (**a**,**c**,**f**). All experiments were repeated three times.

**Extended Data Figure 2.**
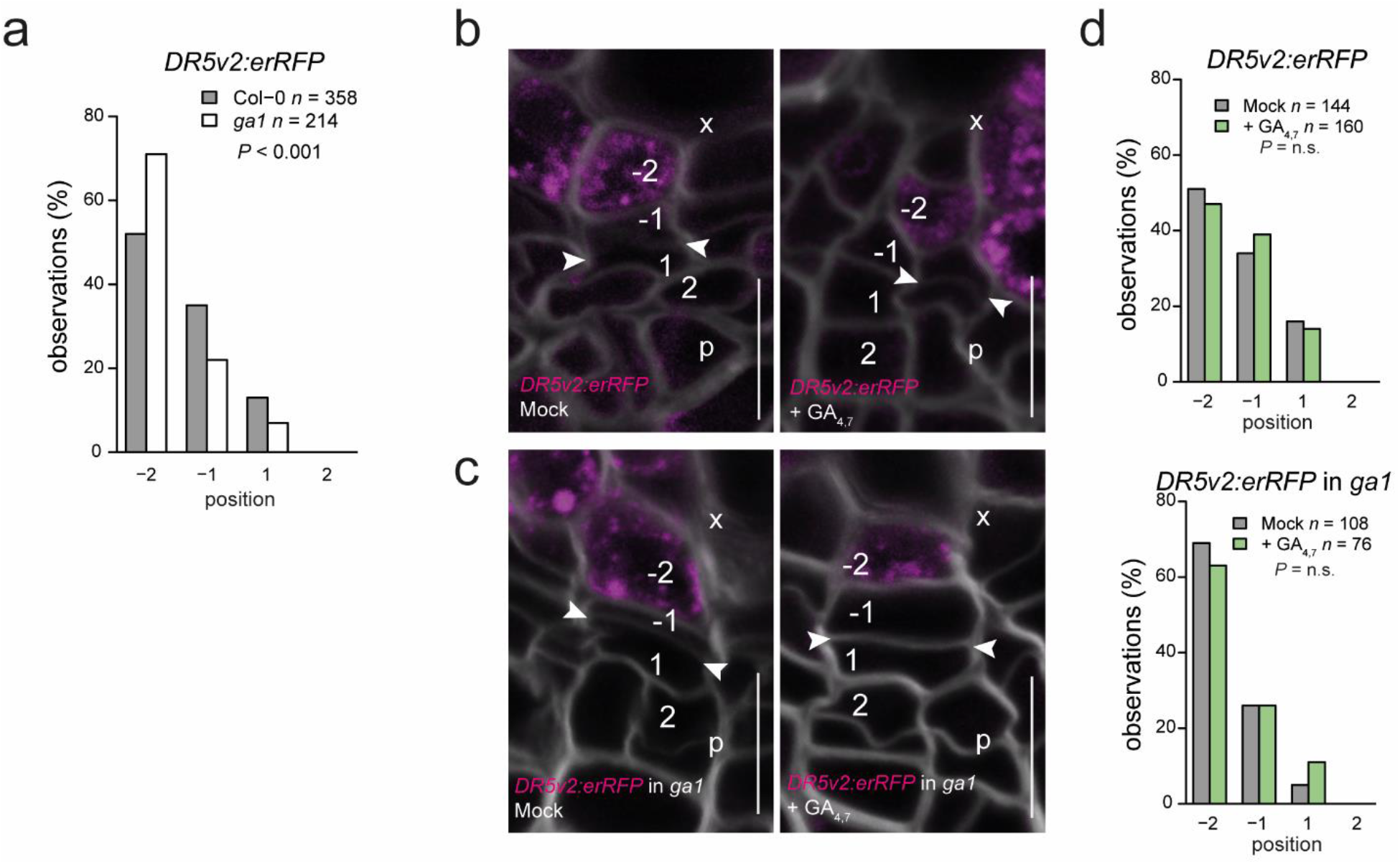
A 24h GA treatment is not sufficient to affect auxin signalling in the cambium. (**a**) Graph comparing the extent of *DR5* in mock-treated 16-day old seedlings of Col-0 and *ga1*. Data is combined from 3 separate repeats. (**b,c**) *DR5v2:erRFP* expression after a 24 h treatment with 2 μM GA4,7 in 14-day old seedlings of Col-0 (**b**) and *ga1* (**c**). The numbers “-2”, “-1”, “1” and “2” indicate the position of the cells relative to the most recent cell division, with negative values towards the xylem and positive towards the phloem. (**d**) Count of the position in the cambium at which the *DR5v2:erRFP* gradient ends. Cellular positions on the x-axis correspond with the cellular positions in panels **b** & **c**. “p”= phloem, “x”= xylem, arrows indicate the most recent cell divisions. Chi-squared test in **a** & **d**. n refers to the total number of observations. All experiments were repeated three times.

**Extended Data Figure 3.**
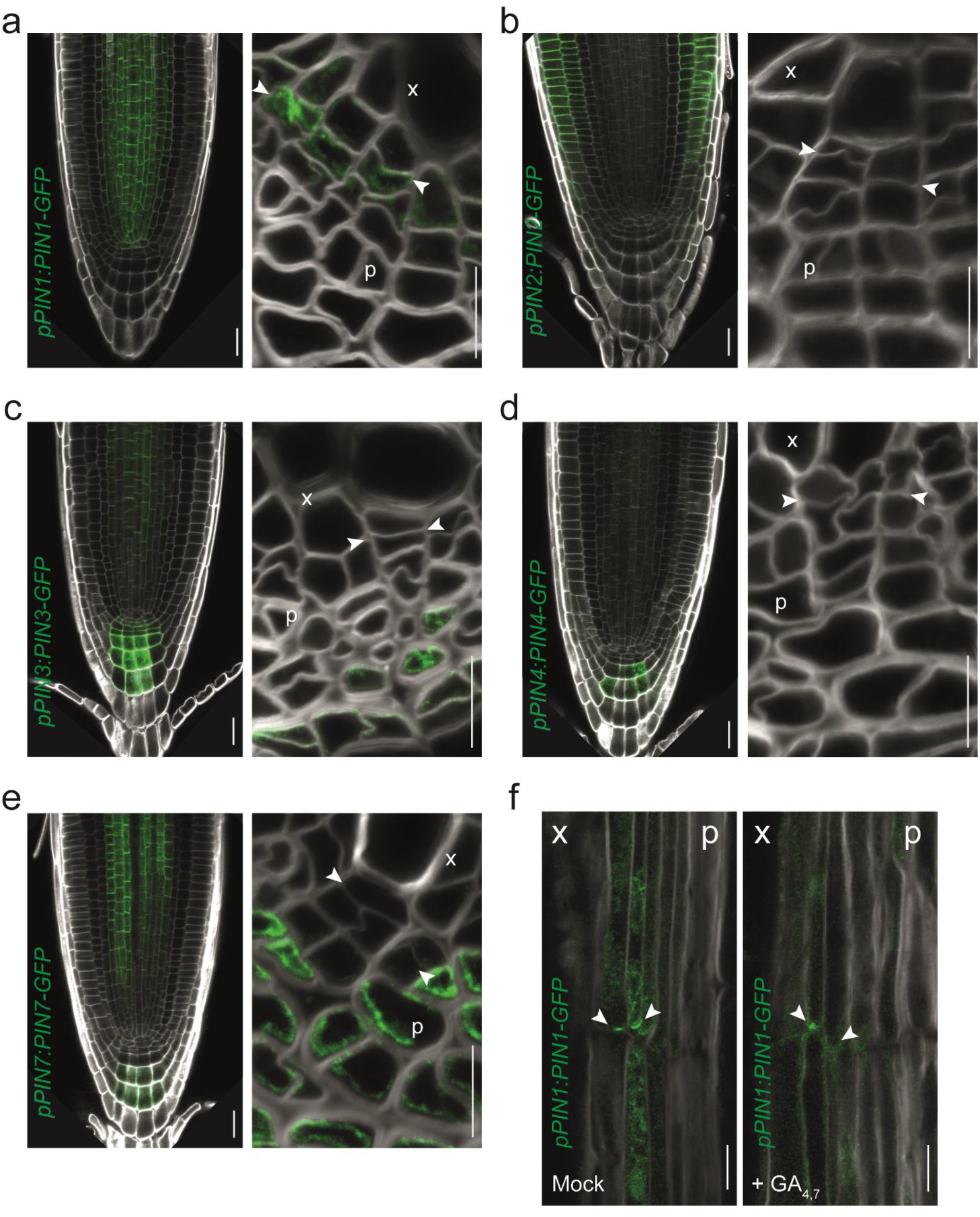
PIN expression patterns in the root tips and vascular cambium. (**a**) pPIN1:PIN1-GFP (**b**) pPIN2:PIN2-GFP (**c**) pPIN3:PIN3-GFP (**d**) pPIN4:PIN4-GFP (**e**) pPIN7:PIN7-GFP expression in 7-day old root tips and 14-day old vascular cambium (**a-e**). Root tips act as positive controls to show that the marker lines have the expected expression pattern in well-studied parts of the root. (**f**) Longitudinal sections showing pPIN1:PIN1-GFP following a 24 h GA or mock treatment in 14-day old plants. “x”= xylem, “p”= phloem. Scale bars are 20 μm in the root tips and 10 μm in the cambium and longitudinal sections. All experiments were repeated three times.

**Extended Data Figure 4.**
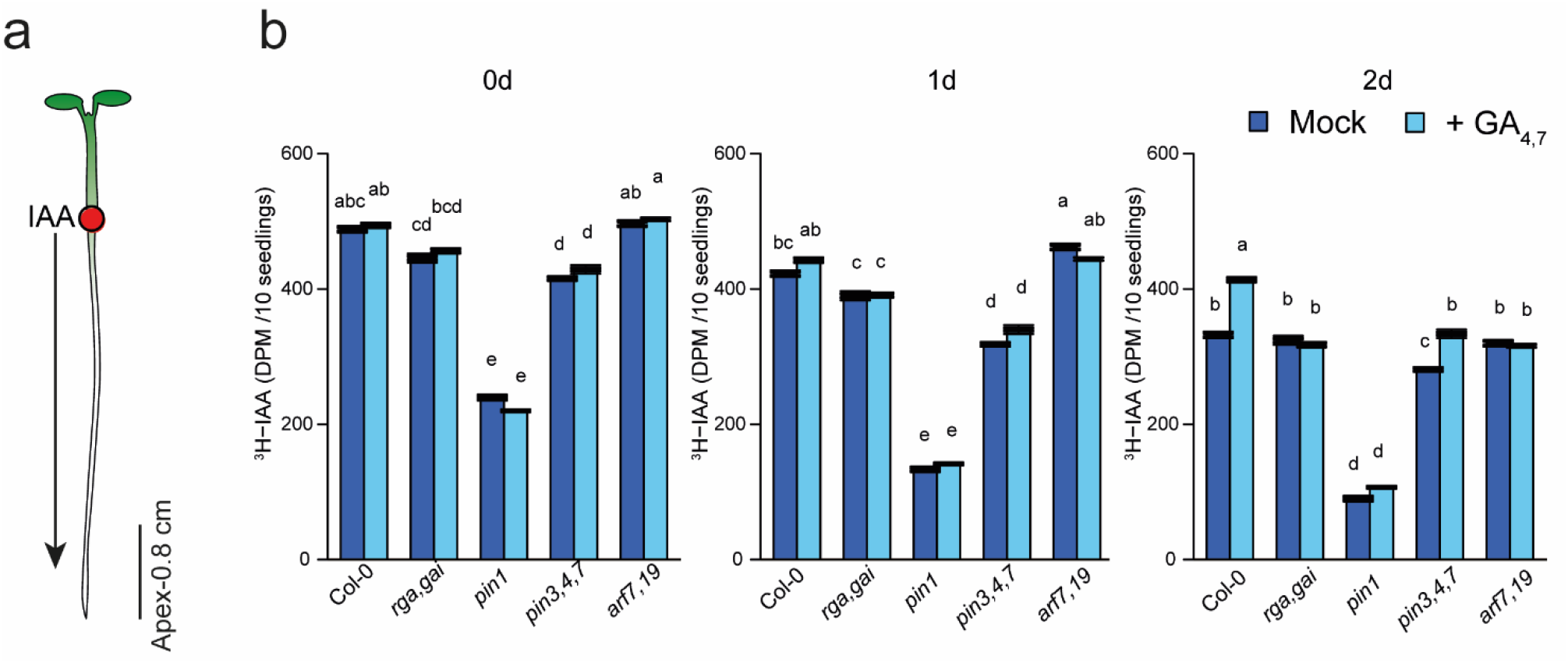
PAT in root tips. (**a**) A schematic explaining the setup of the PAT assay. The red circle indicates the position of ^3^H-IAA application, black arrow showing the direction of IAA movement. The black line marks the area sampled to detect ^3^H-IAA. (**b**) ^3^H-IAA transport from the root-shoot transition zone to the root tips after a 1 h GA treatment in 6-day old seedlings of Col-0 and various mutants. After 0, 1, or 2 days, the plants were treated with ^3^H-IAA for 3 h and then sampled. Data shown are means ±SD (n = 3 independent pools of 10). two-way ANOVA with Tukey’s post hoc test in **b**. Letters indicate a significant difference, with *P* < 0.05.

**Extended Data Figure 5.**
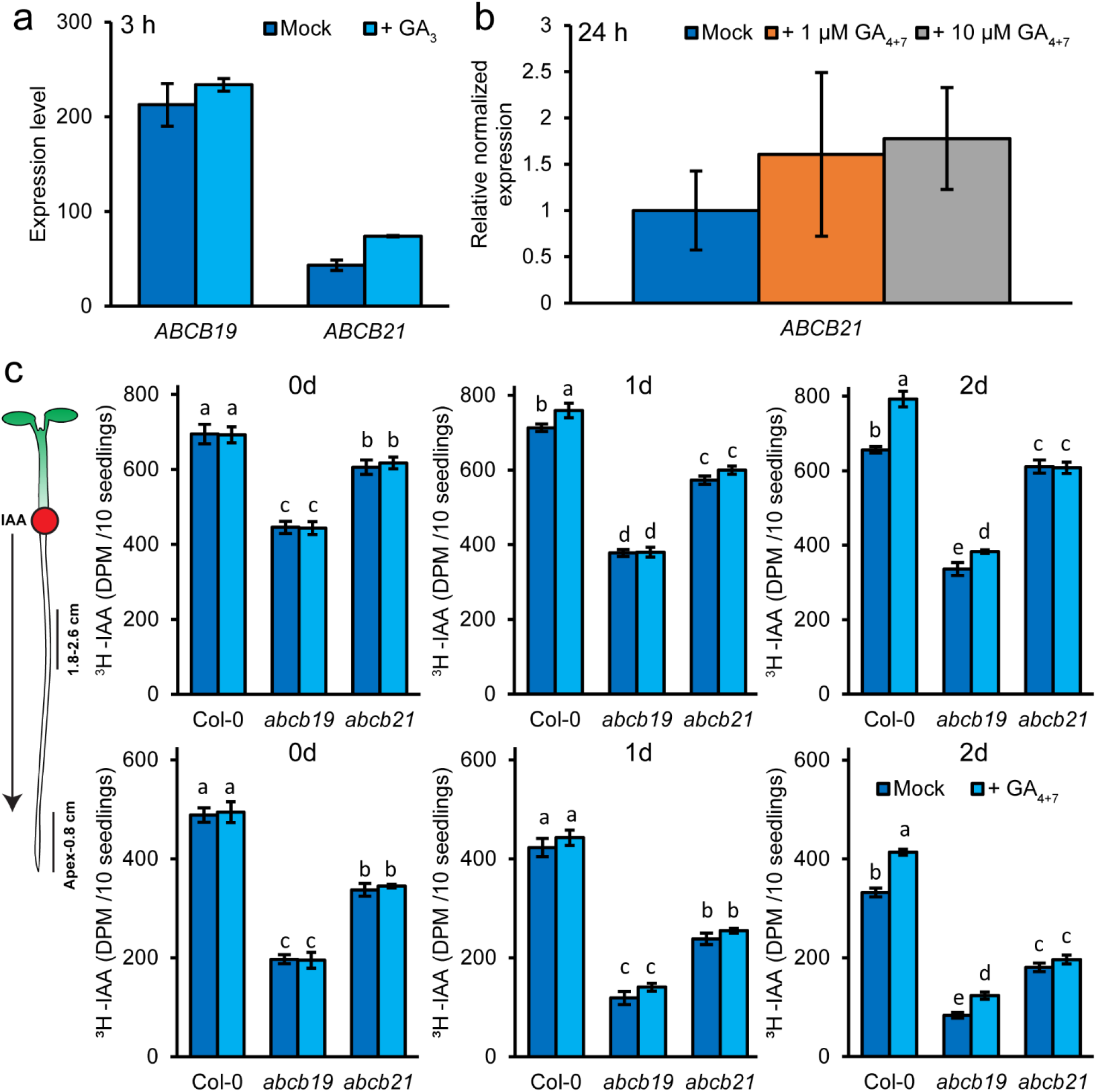
PAT in *abcb* mutants. (**a**) *ABCB19* and *ABCB21* expression 3 h after treatment with 1 μM GA3 from the Arabidopsis eFP Browser^74^. (**b**) Quantitative real-time PCR showing *ABCB21* expression 24 h after treatment with 1 or 10 μM GA_4+7_. Data shown are means ± SD (n = 3 biological replicates, 2 technical replicates). (**c**) ^3^H-IAA transport in *abcb19* and *abcb21* mutant backgrounds from the root-shoot transition zone to 1.8-2.6 mm from the root tips (upper panels) or to the root tips (lower panels) after 1h GA treatment (in 6-days old plants). After 0, 1, or 2 days plants were treated with ^3^H-IAA for 3h and then sampled. ^3^H-IAA transport in Col-0 shown is derived from the same set of experiments shown in Figure 4 and Extended Data Figure 4. Figure Data shown are means ±SD (n = 3 independent pools of 10). two-way ANOVA with Tukey’s post hoc test in **c**. Letters indicate significant difference in p < 0.05.

